# Cytokine response of acute myeloid leukemia cells has impact on patient prognosis: Insights from mathematical modeling

**DOI:** 10.1101/225235

**Authors:** Thomas Stiehl, Anthony D Ho, Anna Marciniak-Czochra

## Abstract

Acute myeloid leukemia (AML) is a heterogeneous disease. One reason for the heterogeneity may originate from inter-individual differences in the responses of leukemic cells to endogenous cytokines. On the basis of mathematical modeling, computer simulations and patient data, we have provided evidence that cytokine-independent leukemic cell proliferation is linked to early relapses and poor overall survival. Depending whether the model of cytokine-dependent or cytokine-independent leukemic cell proliferation fits to the clinical data, patients can be assigned to two groups that differ significantly with respect to overall survival. The modeling approach further enables us to identify parameter constellations that can explain unexpected responses of some patients to external cytokines such as blast crisis or remission without chemotherapy.

## Introduction

Acute myeloid leukemias (AML) comprise a heterogeneous group of malignant diseases. Since major clinical symptoms originate from impairment of healthy blood cell production, it is important to understand how leukemic cells interfere with healthy hematopoiesis. Clinical and genetic observations reveal a strong heterogeneity among individual patients. One reason for the observed heterogeneity may be differences in cytokine dependence of leukemic cells, i.e., cells of some patients require cytokines to expand (cytokine-dependent leukemic cells) whereas others exhibit autonomous (cytokine-independent) growth.

The idea that cytokine dependence of leukemic cells differs between patients is supported by experimental results. Xenotransplantation assays reveal that some leukemia samples exclusively engraft in mice transgenic for human cytokines and not in standard NSG mice.^1,2^ Similarly, in vitro studies imply that leukemic cells of some patients exhibit autonomous growth in cell cultures whereas others require cytokines to expand.^3–5^ The correlation between cytokine-dependence in cell culture and patient survival suggests that cytokine dependence of leukemic cells may be a clinically meaningful parameter.^6^ However, it can depend on the culture conditions whether a leukemia sample exhibits autonomous growth or not.^3^

Cytokine administration has become a widely used supportive strategy to prevent chemotherapy-related neutropenia.^7^ In this context the question arises whether cytokines could potentially stimulate leukemic cells that survived therapy and trigger relapse. Although studies in AML patients suggest that leukemic cells can be recruited into cell cycle in response to administered cytokines,^8–10^ multiple clinical trials imply that supportive cytokine treatment has no negative effects on relapse free survival.^7^ Nevertheless, there exist trials and case reports stating that in some patients administration of cytokines or their analogues increases leukemic cell load or reduces relapse free survival.^11–13^ Different genetic hits accounting for that have been identified so far.^12,14,15,16^ On the other hand, there exist reports of patients achieving complete remission solely by cytokine administration without chemotherapy.^17–25^ Both phenomena, negative and positive impact of cytokines on leukemic cell load, are so far not well understood.

The aim of this work is to study if cytokine dependence of leukemic cells has an impact on the clinical course of the disease. For this purpose, we compare disease dynamics in case of cytokine-dependent (i.e. leukemic cells require endogenous cytokines to expand) and cytokine-independent (i.e. leukemic cells can expand in absence of endogenous cytokines) AMLs using mathematical models. We focus on the following questions: (i) How does time evolution of blasts differ in cytokine-dependent and cytokine-independent AML? Can we conclude from bone marrow aspiration data of relapsing acute myeloid leukemia patients whether leukemic cells require cytokines to expand? (ii) Does it have a prognostic impact if patient data fits to the model of cytokine-dependent or to the model of cytokine-independent AML? (iii) Which cell parameters determine whether cytokine administration has negative, neutral or positive effects on the leukemic cell load?

To approach these questions, we develop new mathematical models of cytokine-dependent and cytokine-independent AML and apply them to patient data showing time changes of bone marrow blast counts between first remission and relapse. Comparing the two models we identify key dynamic features that help to distinguish between both scenarios. Model-based patient data analysis suggests that the overall survival depends on the type of regulatory feedback governing cancer stem cell behavior and it is significantly worse in case of cytokine-independent AML. Mathematical models allow also to explain the unexpected response of patients to cytokines described in literature.^11–13,17–22^

Mathematical models are a useful tool to understand processes that cannot be manipulated or measured experimentally. They allow rigorous comparison of different hypothetical scenarios and estimation of unknown parameters.^26^ Studies from literature demonstrate that mathematical modeling is a suitable approach to investigate the dynamics of cancer cells subjected to regulatory feedbacks or treatment interventions.^26–30^ Especially in case of ambiguous experimental results or in systems where the observables strongly depend on experimental conditions, a model-based interpretation of patient data can provide additional insights.

## Methods and Model Description

### Mathematical models

The interaction of healthy and leukemic cells by cytokine feedbacks and consumption of environmental resources, such as niche spaces, makes it necessary to model both healthy and leukemic cells. To study the impact of these interactions on the clinical course of the disease our models incorporate two different modes of feedback, namely cytokine-dependent leukemic cells and cytokine-independent leukemic cells. Cytokine-dependent leukemic cells expand only in presence of cytokines, whereas cytokine-independent (autonomous) leukemic cells have the ability to expand without cytokine stimulation. In this work we develop a new mathematical model for cytokine-independent leukemic cells and compare it to a model of cytokine-dependent leukemic cells proposed by Stiehl et al^31^ and applied to data from AML patients.^26^ Both models are an extension of a model of hematopoiesis^32^, which has been validated on the basis of patient data and applied to clinical questions.^26,33,34^

In the following we introduce the biological system underlying our models. In a multi-step process, the hematopoietic stem cell (HSC) population gives rise to all types of mature blood cells.^35^ Since acute myeloid leukemia (AML) is a disease of the myeloid lineage, our models focus on the granulopoietic branch of the hematopoietic system. A complex network of cytokine signals adjusts cell production to the need of the organism.^36^ Time evolution of the non-linear cytokine feedback in the models are inspired by G-CSF, the main cytokine of granulopoiesis,^36^ and were proposed by Marciniak-Czochra et al.^32^ It has been reported that cytokine concentration influences properties of stem cells^37^ and more mature cells,^38^ we therefore assume in the model that feedback signals act on all mitotic cell compartments.

In case of AML the leukemic cell population shows a similar hierarchical organization as the hematopoietic system with the leukemic stem cell (leukemia stem cell, LSC, leukemia initiating cell, LIC) population at the top of the hierarchy.^39,40^ To investigate the impact of leukemic cell cytokine dependence on disease dynamics we consider two models, which are summarized in Figure 1 and Table 1.

**Figure 1:**
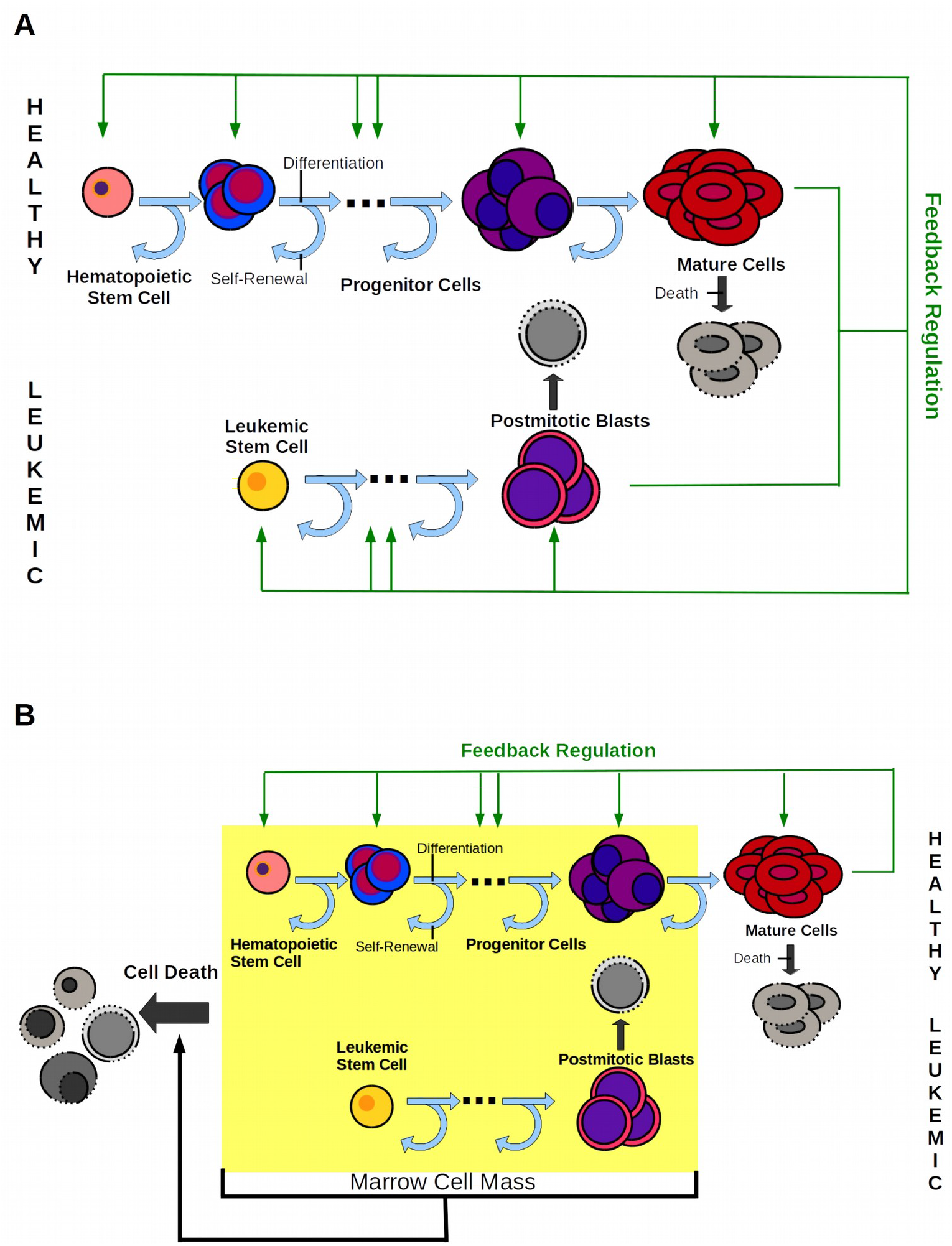
Models of cytokine-dependent and cytokine-independent AML. (A) Model 1: Hematopoietic and leukemic cells depend on the same cytokine. (B) Model 2: The leukemic cells are independent of cytokines. Crowding in marrow space results in increased apoptosis. The fraction of self-renewal assigned to non-stem cells is a measure of the average number of cell divisions performed before a cell becomes post-mitotic under homeostatic conditions.^33^

**Table 1:**
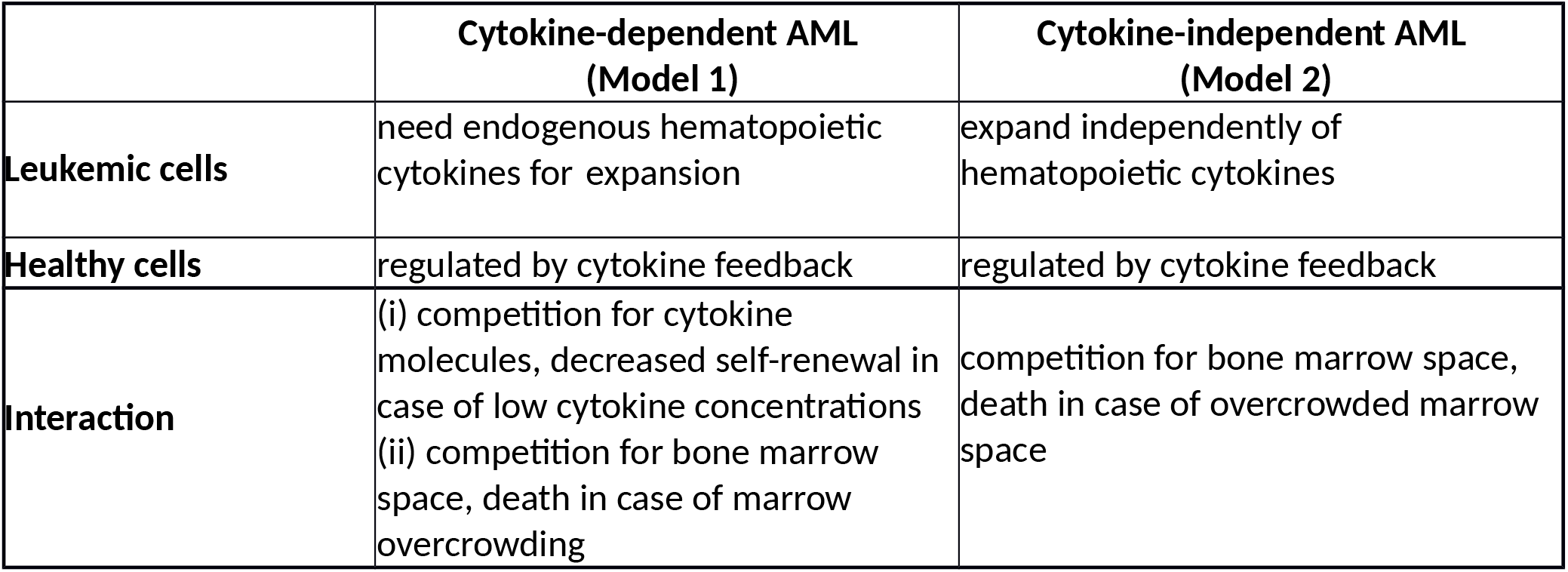
Comparison of AML models

|  | **Cytokine-dependent AML**<br>**(Model 1)** | **Cytokine-independent AML**<br>**(Model 2)** |
| --- | --- | --- |
| **Leukemic cells** | need endogenous hematopoietic cytokines for expansion | expand independently of hematopoietic cytokines |
| **Healthy cells** | regulated by cytokine feedback | regulated by cytokine feedback |
| **Interaction** | (i) competition for cytokine molecules, decreased self-renewal in case of low cytokine concentrations<br>(ii) competition for bone marrow space, death in case of marrow overcrowding | competition for bone marrow space, death in case of overcrowded marrow space |

In Model 1 (cytokine-dependent AML), it is assumed that leukemic cells depend on the same endogenous cytokines as healthy hematopoietic cells. This assumption is justified by the following biological findings: (1) Leukemic cells express the same cytokine receptors as hematopoietic cells.^41,42^ (2) Leukemic cells of some patients expand only in the presence of cytokines and engraft only in mice transgenic for human cytokines.^2,4^ (3) Cytokine administration in AML patients recruits leukemic cells into S-phase.^8–10^ Since hematopoietic and leukemic cells absorb and degrade cytokine molecules by receptor-mediated endocytosis, the two cell lineages interact through competition for the cytokine.^36,41,42^

Model 2 (cytokine-independent AML) is based on the evidence that in some patients malignant cells are autonomous with respect to physiological hematopoietic growth factors.^6,43–45^ The cytokine-independent leukemic cell growth is then limited by a competition for the bone marrow space that results in an increased cellular degradation due to overcrowded bone marrow space. It is modeled by a feedback loop in death rates depending on the total immature cell population. A model assuming increased cell differentiation in case of marrow overcrowding leads to similar results. These assumptions are justified by the following evidence: (1) Different mechanisms of cytokine-independent leukemic cell expansion have been identified, such as autocrine stimulation or constitutive activation of key signaling pathways due to mutations.^43,44,46^ (2) Markers for cell death such as LDH, are increased in bloodstream of leukemic patients^47–50^ and enhanced cell death is observed in marrow samples of many patients.^51^ The increased cell death is included in Model 2. Several mechanisms for spatial competition have been described such as (i) physical stress owing to overcrowding leading to extinction of cells,^52^ (ii) competition for a limited niche surface expressing certain receptors (contact molecules) necessary for survival of healthy and leukemic cells^53,54^ and apoptosis (cell death) occurring, if no contacts to these molecules can be established.^55^

The latter niche competition mechanism is also valid for the cytokine-dependent AML in case of overcrowded bone marrow. Therefore, we include it also in the Model 1. As shown by numerical simulations this mechanism is relevant in Model 1 only under external cytokine stimulation. In absence of external cytokine stimulation, the feedback mechanism prevents significant overcrowding of the bone marrow space in cytokine-dependent AMLs.

Both considered models include one leukemic cell lineage and one healthy cell lineage. Dynamics of the different healthy and leukemic cell types are given by ordinary differential equations. We assume that each lineage can be described by 2 compartments: dividing cells (including stem cells and progenitors) and post-mitotic cells. The resulting four-compartment model architecture is based on a simplified description of the multi-stages differentiation process, which reduces the complexity of the differentiation process to focus on mechanisms and effects of competition between different cell lines. The previous studies showed that reduction of the number of compartments do not change the behavior of the model after the modification of the parameters corresponding to the new interpretation of the compartments.^26^

Each cell type is characterized by the following cell parameters, which are summarized in Figure 2:

- *Proliferation rate*, describing the frequency of cell divisions per unit of time.
- *Fraction of self-renewal*, describing the fraction of progeny cells returning to the compartment occupied by the parent cells that gave rise to them. On the basis of our earlier work and on compatibility with clinical data,^26,32,34^ we assume that the fraction of self-renewal is regulated by feedback signaling. The fraction of self-renewal assigned to non-stem cells is a measure of the average number of cell divisions performed before a cell becomes post-mitotic under homeostatic conditions.^33^
- *Death rate*, describing the fraction of cells dying per unit of time. For simplicity, we assume that under physiological conditions dividing cells do not die and non-dividing cells die at constant rates.

**Figure 2:**
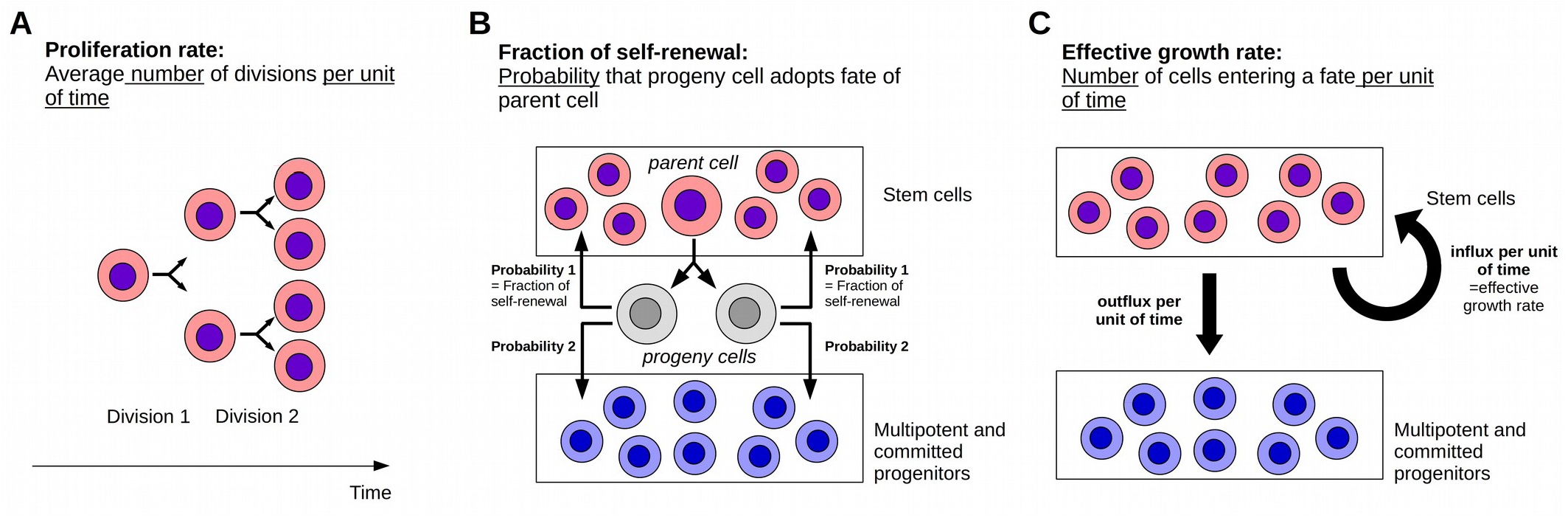
Important model parameters. (A) The proliferation rate describes the average number of divisions performed by a cell of given type. (B) The fraction of self-renewal describes the probability that a progeny cell originating from division adopts the fate of its parent cell, e.g., that a daughter cell of a stem cell is again a stem cell. (C) The effective growth rate describes how many cells enter a given cell fate per unit of time, e.g., the number of stem cells that are generated per unit of time. The effective growth rate depends on the proliferation rate and the fraction of self-renewal.

For analysis of the model dynamics, we define also the *effective growth rate* of a mitotic cell population. It describes how many mitotic cells are generated per unit of time, see Figure 2. This quantity depends on the proliferation rate and the fraction of self-renewal of the immature cell population. A high proliferation rate and a low fraction of self-renewal, as long as it is larger than one half, can lead to the same effective growth rate as a combination of high self-renewal and slow proliferation. The supplementary information contains model details (Section 1) together with analytical studies (Section 2) and model parametrization (Section 3).

### Simulations

Simulations have been performed using standard ODE-solvers in MATLAB (Version 7.8, The MathWorks, Inc.). We start simulations with equilibrium cell counts in the hematopoietic lineage and a small number of LSC (1 per kg of body weight), mimicking the appearance of LSC due to a mutation or survival of LSC after therapy. Parameters for equations of the healthy cell line are taken from the previous work^34^ (see supplementary information Section 3.2). Leukemic cell parameters are selected from biologically plausible ranges to fit the patient data using the least square method (supplementary information Section 3). The fitting procedure is used to determine which of the two models fits to the data. During the fitting process we allow proliferation rates to vary between one division per two years and one division per day. The latter is considered the upper bound taking into account the DNA replication time of a eukaryotic cell.^26^ Self-renewal may vary between zero and one. Blast half-life is chosen between 25% and 100% of leukocyte half-life, motivated by literature.^56^

To decide which of the models fits to the data of a given patient better, we find the best fit for each of the two models and calculate the RMSE (root mean square deviation) for each model. If RMSE for Model 1 is 5% larger than that for Model 2, we conclude that Model 2 fits the data better than Model 1 and vice versa. The results are robust with respect to these choices.

### Application to patient data

We use bone marrow aspiration data from patients participating in clinical trials at the University Hospital of Heidelberg (Department of Medicine V; Heidelberg, Germany). Written consent for usage of clinical data for scientific purposes was obtained from each patient. We consider the data of 41 randomly chosen patients. Patients meet the following criteria: (i) at least one documented relapse of the disease in the bone marrow, (ii) achievement of complete hematological remission after treatment of primary diagnosis, (iii) successful bone marrow examination at relapse, and (iv) documented date of death or patients were still alive at the day of data collection. Criterion (iv) limited the number of considered patients.

### Statistical analysis

Survival distributions of different patient groups are compared using standard Kaplan-Meier survival analysis.^57^ In all considered cases, the test yields significant results (p < 0.05).

### Data availability

All data analyzed in this study are included in this article, its supplementary information and in the cited references.

## Results

### Cytokine-dependence of leukemic cells has an impact on blast expansion rates

To study how the dependence of leukemic cells on endogenous cytokines impacts on the clinical course of AML, we introduce two different mathematical models. In one model leukemic cells need endogenous cytokines to expand (cytokine-dependent AML) whereas in the other model leukemic cells can expand independently of endogenous cytokines (cytokine-independent AML). The models are summarized in Figure 1 and in Table 1. They are extensions of previous works and have been parameterized based on patient data.^26, 31-35^ For details see section “Methods and Model Description” and supplementary information Sections 1 and 3.

We perform numerical simulations to compare blast expansion rates in case of cytokine-dependent and cytokine-independent (autonomous) leukemic cells. To simulate the fastest possible expansion of leukemic cells, we set their proliferation rate to one division per day, which is an upper bound.^58^ We then measure the time span from origin of one LSC per kg of body weight to a marrow blast fraction of 10% for different LSC self-renewal fractions, see Figure 3. The simulations show that, for a given leukemic cell self-renewal, leukemic cell expansion is faster for cytokine-independent leukemic cells compared to cytokine-dependent leukemic cells. Whereas for cytokine-dependent leukemic cells a minimum of 200 days is needed to obtain a marrow blast fraction of 10%, 30 days are sufficient in case of cytokine-independent leukemic cell expansion. This observation can serve as a discriminator between both models and suggests that fast relapses can be explained only by autonomous leukemic cell growth or treatment failure. The latter can be excluded based on bone marrow biopsy or aspiration.

**Figure 3:**
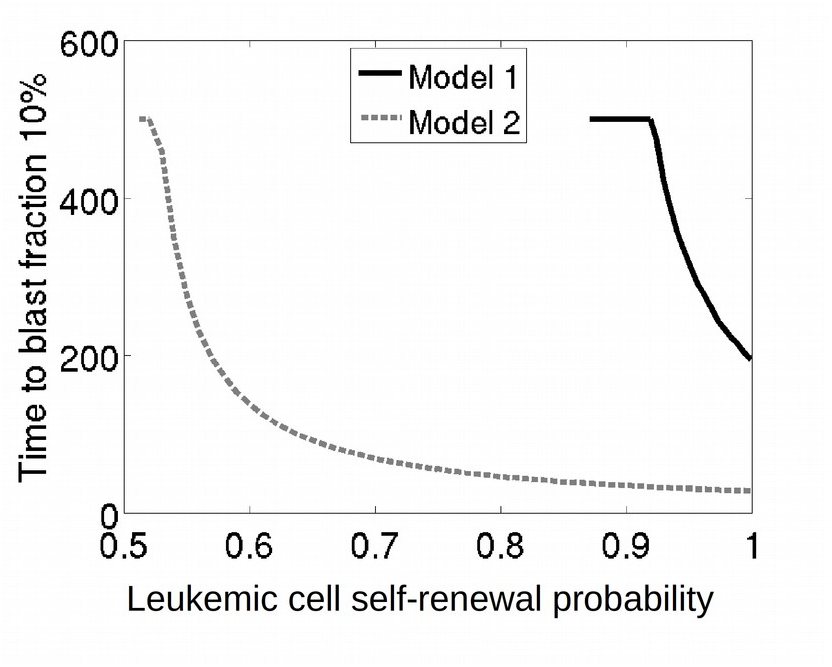
Leukemic cell dynamics in Model 1 and Model 2. At *t*=0 one leukemic cell per kg of body weight is added to the healthy equilibrium. The time elapsing until a marrow blast fraction of 10% is depicted depending on leukemic cell self-renewal. The simulations imply that leukemic cell expansion is faster in Model 2 (dashed line) compared to Model 1 (solid line).

### Cytokine-dependence of leukemic cells contributes to inter-individual heterogeneity of clinical courses

To check whether the observed clinical courses are covered by our models, we fit both models to patient blast counts between complete remission (CR) and relapse, see Figure 4. Time evolution of blast counts of 22 of the considered 41 patients are compatible with both models (Figure 4A). Evolution of blast counts of 17 patients are compatible only with the model of cytokine-independent AML (Figure 4B) and blast counts of 2 patients are compatible only with the model of cytokine-dependent AML (Figure 4C). The fitting results demonstrate that our models capture clinical data and that cytokine dependence of leukemic cells can contribute to the observed clinical heterogeneity of acute myeloid leukemias.

**Figure 4:**
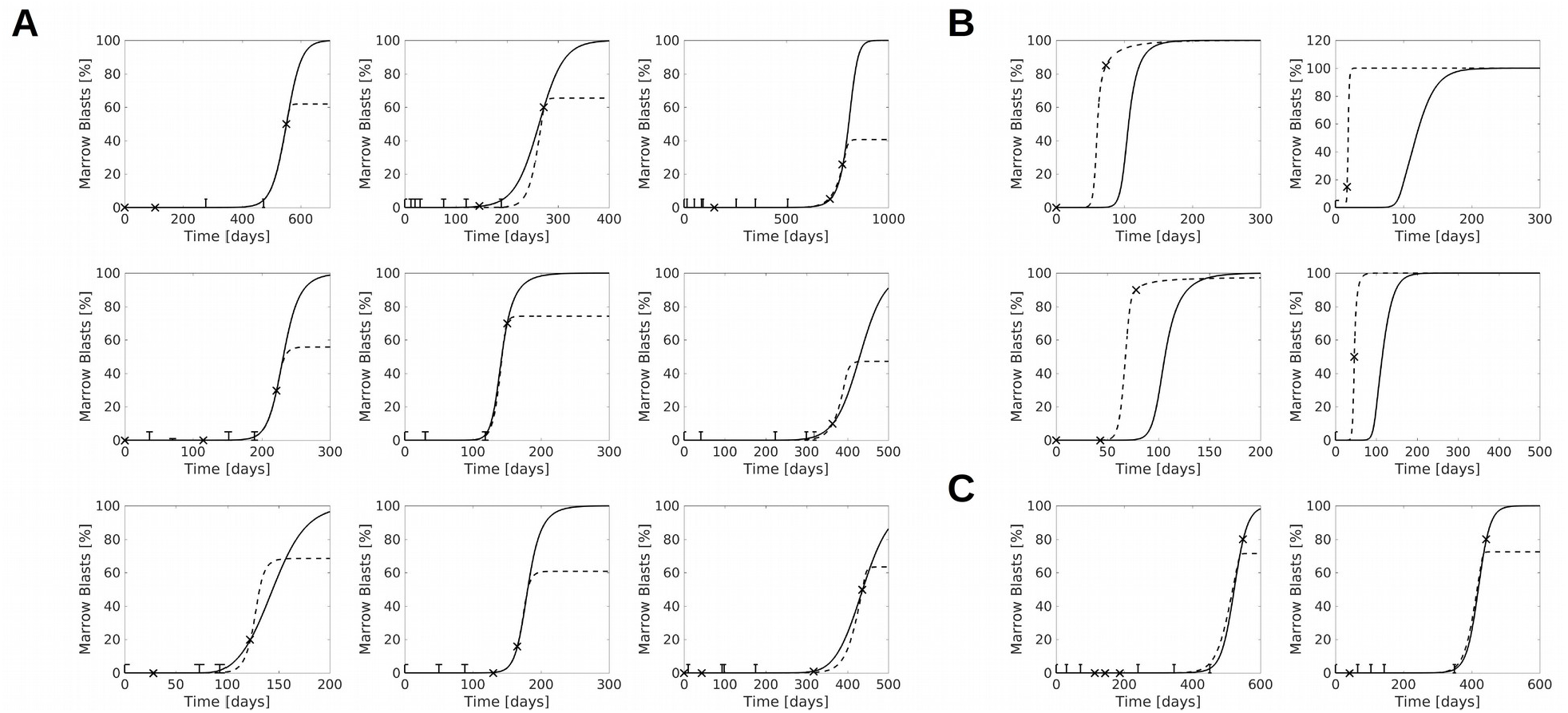
Fit of models to patient data. Models are fitted to marrow blast fractions of AML patients between remission and relapse (solid line: Model 1; dashed line: Model 2). Bone marrow blast fractions are marked by “x” if exact values were reported, if intervals, such as “less than 5% blasts” have been reported, the data is shown as a bar. Panels: Examples for patient data compatible with Models 1 and 2 (A), Model 2 but not Model 1 (B), or Model 1 but not Model 2 (C).

### Prognostic significance of autonomous leukemic cell growth

We ask whether cytokine-dependence of leukemic cells has impacts on patient prognosis. For this purpose, we compared overall survival of patients compatible only with the model of cytokine-independent AML (group 2, n=17) to patients compatible with the model of cytokine-dependent AML (group 1, n=24). Survival analysis yields a significantly longer overall survival for group 1 (p=7 × 10^−3^, Figure 5, median overall survival 700 days) compared to group 2 (median overall survival 350 days). The survival after the first relapse is also significantly increased for group 1 (p=0.03). These results are in line with reports from literature stating that autonomous leukemic cell growth has a negative impact on patient prognosis.^6^

**Figure 5:**
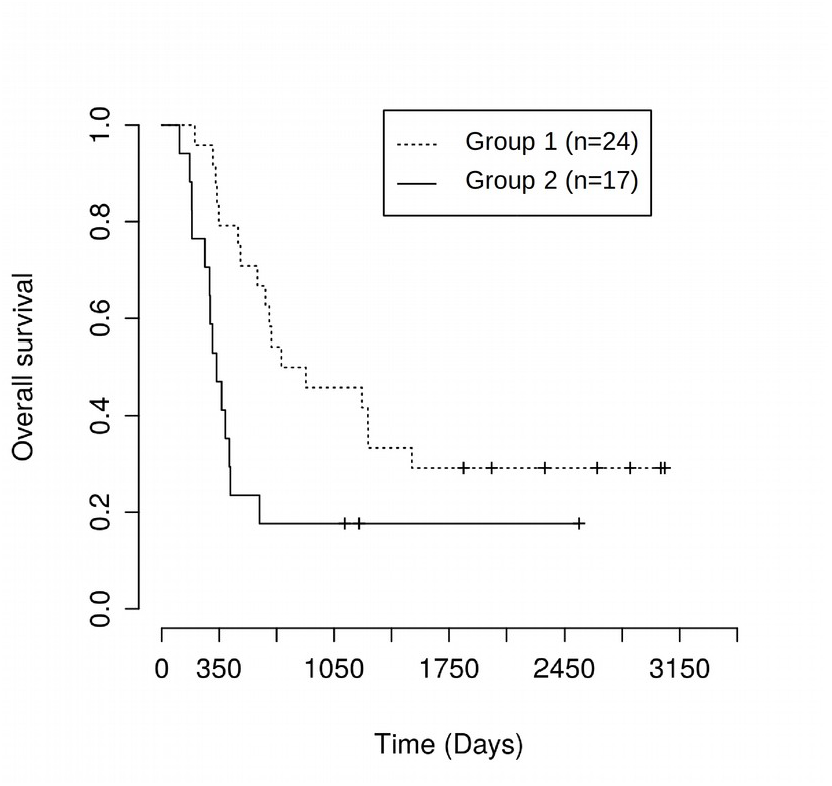
Patient data compatible only with model of cytokine independent AML correlates with poor overall survival. Overall survival of two patient groups is compared. Group 1: Patients compatible with model of cytokine dependent AML, group 2: Patients compatible with model of cytokine independent AML and incompatible with model of cytokine dependent AML. Survival of both groups differs significantly (p=7 × 10^−3^). Similar results are obtained if survival after first relapse is compared (p=0.03).

### Potential effects of supportive cytokine administration

Depending on the underlying leukemic cell properties and feedbacks, external cytokine administration may have divergent effects. In some patients, external cytokines stimulate blast expansion at the expense of healthy cells,^11^ whereas in other patients leukemic cells are out-competed by stimulated healthy cells which results in a reduction of blast counts.^18^ To understand which cell parameters are crucial for the observed dynamics we run model simulations for multiple parameter combinations. Representative results are depicted in Figure 6. The results can be classified using the effective growth rate of mitotic healthy and leukemic cells. It describes how many mitotic cells of the respective type are produced per unit of time (Figure 2). In case of cytokine-dependent AML, cytokine administration is beneficial if the effective growth rate of immature leukemic cells is smaller than the corresponding rate for the healthy cells (Figure 6A) and harmful otherwise (Figure 6B). Note that in case of cytokine-dependent AML a higher self-renewal of leukemic cells compared to hematopoietic cells is sufficient for leukemic cell expansion, independent of the relation between the corresponding effective growth rates. If the effective growth rates of healthy and leukemic cells are similar, cytokine administration has no relevant effect. The simulation depicted in Figure 6A is in line with the observation that in some patients one cycle of cytokine administration can result in complete remissions that are stable for time intervals between several months and more than one year.^18,20,21^ The simulation depicted in Figure 6B is in line with the observation that cytokine stimulation can result in blast crisis.^11^ A fit of our models to the data from Duval et al^11^ is depicted in Figure 7.

**Figure 6:**
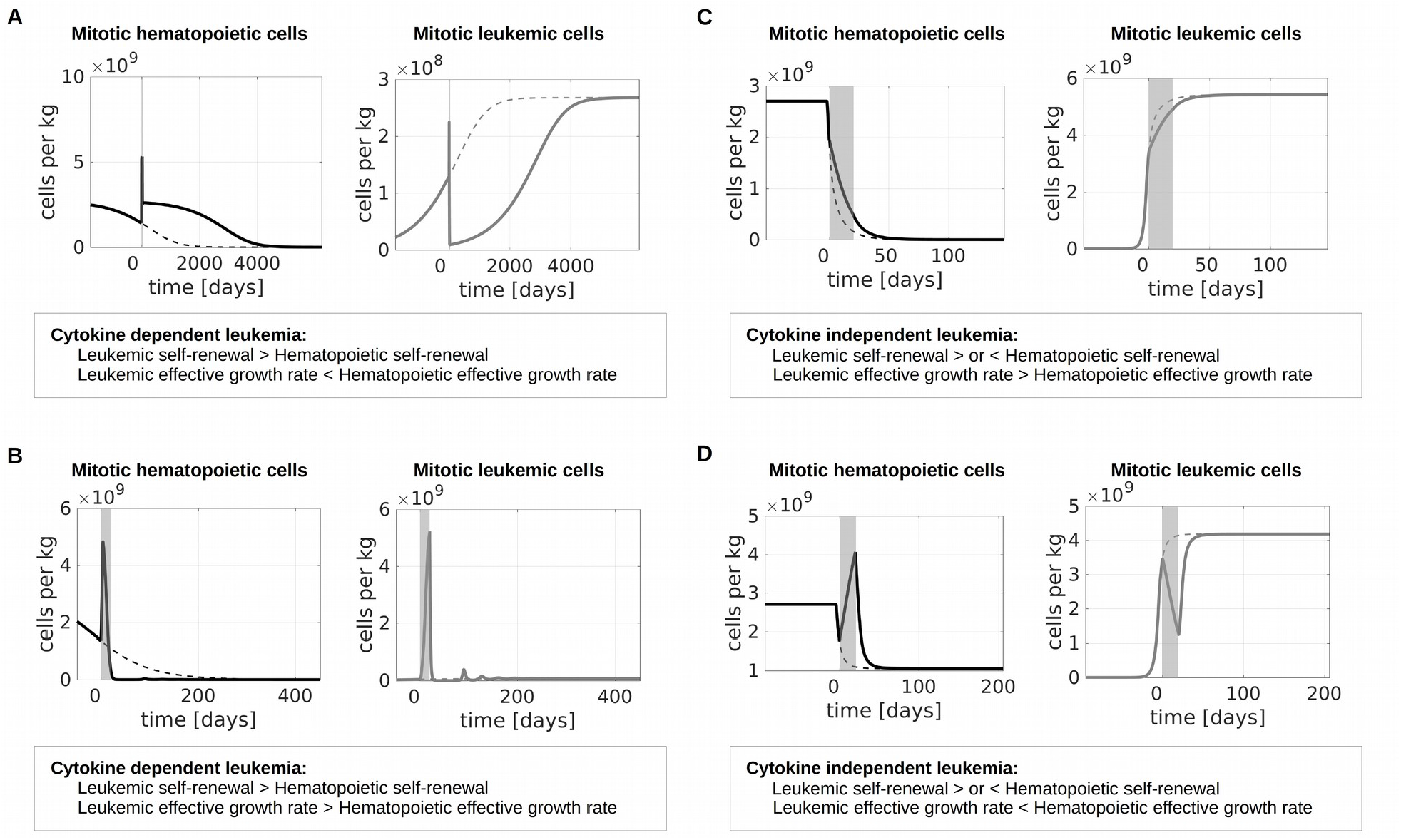
Effect of cytokine stimulation on leukemic cell burden. When mature cell counts are reduced by 50% due to leukemic cell load we simulate cytokine stimulation for 20 days. Dotted lines show dynamics in absence of cytokine stimulation, solid lines show dynamics during and after cytokine stimulation. (A) Cytokine dependent AML, the self-renewal of leukemic cells is higher than the self-renewal of hematopoietic cells and the effective growth rate of leukemic cells is smaller than the effective growth rate of hematopoietic cells. Cytokine administration reduces the leukemic cell burden. (B) Cytokine dependent AML, the self-renewal of leukemic cells is higher than the self-renewal of hematopoietic cells and the effective growth rate of leukemic cells is higher than the effective growth rate of hematopoietic cells. Cytokine administration increases the leukemic cell burden. (C) Cytokine independent AML, the effective growth rate of leukemic cells is larger than the effective growth rate of hematopoietic cells, leukemic cell self-renewal can be larger or smaller than hematopoietic cell self-renewal. Cytokines cannot reduce the leukemic cell burden, but they slightly reduce leukemic cell expansion. (D) Cytokine independent AML, the effective growth rate of leukemic cells is smaller than the effective growth rate of hematopoietic cells, leukemic cell self-renewal can be larger or smaller than hematopoietic cell self-renewal. In this case leukemic cell counts can be reduced by cytokine administration.

**Figure 7:**
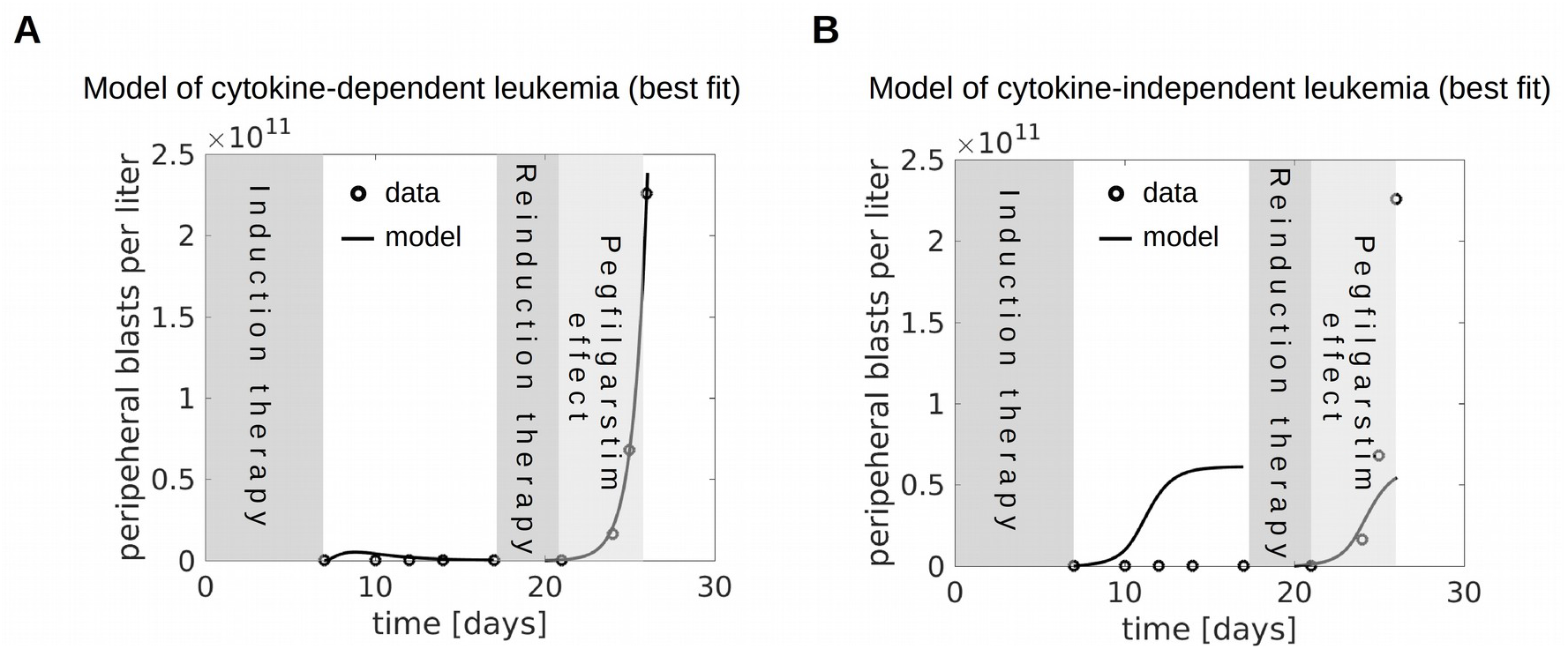
Blast crisis after pegfilgrastim. Blast crisis after pegfilgrastim administration can only be explained by cytokine sensitive leukemic cells. The figure shows the best fits of the models of cytokine-dependent (A) and cytokine independent (B) AML to the data from Duval.^11^

Also in the model of cytokine-independent AML the effect of cytokine administration depends on the effective growth rate of mitotic leukemic and hematopoietic cells. If the effective growth rate of leukemic cells is larger than that of hematopoietic cells, cytokine administration cannot reduce leukemic cell load, it only leads to a negligible reduction of the speed of leukemic cell expansion (Figure 6C). In the opposite scenario cytokine administration can temporarily reduce the leukemic cell burden but remission, if achieved at all, lasts shorter than in the case of cytokine-dependent AML. The scenario depicted in Figure 6D fits to the clinical observations reported by Xavier et al.^20^ Repeated cytokine administration can be used in this case to control the leukemic cell burden.^20^

## Discussion

In this work we compare the clinical course of cytokine-dependent and cytokine-independent acute myeloid leukemias. We focus on two scenarios. In Scenario 1 (cytokine-dependent AML) leukemic and healthy cells compete for endogenous cytokines, which they require for expansion. In Scenario 2 (cytokine-independent AML) leukemic cells expand independently of cytokines but cells compete for the bone marrow space and die in case of overcrowding. Both scenarios are supported by experimental evidence.^3–5,43,46^

Unlike many previous models, the models considered in this work explicitly include dynamics of healthy cells. Animal experiments show that hematopoietic stem cells isolated from leukemic organisms can lead to normal hematopoiesis if transplanted to a non-leukemic host.^59^ This finding suggests that a better understanding of the disease mechanisms could allow to derive clinical strategies that support healthy hematopoiesis and reduce expansion of leukemic cells.

The proposed mathematical models provide criteria allowing discrimination between cytokine-dependent and cytokine-independent AMLs. Our results suggest that rapid leukemic cell expansion and early relapses are typical for cytokine-independent AML. The models suggest that if time between complete remission and 10% marrow blasts at relapse is shorter than 200 days, only the model of cytokine-independent AML is able to explain the dynamics.

Fitting of the developed models to patient blast counts between complete remission and relapse demonstrates that patients compatible only with the model of cytokine-independent AML have a significantly poorer overall survival compared to patients compatible with the model of cytokine-dependent AML. This observation is in line with data from cell culture studies showing that autonomous cell growth is correlated with a poor prognosis.^6^

Another important difference between the models lies in the reaction of the patient to external cytokine administration. Cytokine administration is a commonly used treatment strategy to increase healthy cell counts and to reduce complications of chemotherapy.^7^ Although considered safe in general, there exist multiple reports of patients showing unexpected increase or reduction of the leukemic cell burden after cytokine administration. Increasing leukemic cell counts can be explained by cytokine-mediated blast expansion,^11^ whereas decreasing leukemic cell counts result from stimulation of hematopoiesis and out-competition of leukemic clones.^18^ Our modeling approach identifies the effective growth rate of leukemic and hematopoietic mitotic/stem cells as a crucial parameter to understand divergent reaction of patients to cytokine administration. The effective growth rate of a cell population describes how many cells of that population are produced per unit of time. It can potentially be estimated based on mathematical models. Our simulations show that cytokine stimulation can induce complete remissions even in cytokine-dependent AML provided the effective growth rate of mitotic leukemic cells is smaller than that of mitotic hematopoietic cells (note that an effective growth rate of leukemic cells larger than zero is necessary and sufficient for their expansion). This scenario is in line with multiple clinical observations.^17–19,21,22^ If the effective growth rate of leukemic cells is higher compared to hematopoietic cells, cytokine administration can result in blast expansion as it has been clinically observed.^11^

Similarly, leukemic cell load can be temporarily reduced in cytokine-independent AML if the effective growth rate of hematopoietic cells is larger than that of leukemic cells. Nevertheless, this reduction is less efficient compared to the case of cytokine-dependent AML, since remaining leukemic cells expand fast. Model dynamics in this case also fit to clinical reports.^20^ The observation that a given treatment can be beneficial in some patients and harmful in others demonstrates the need to distinguish between the regulation modes present in different patients. The criteria developed in this work can be helpful for this task.

The models considered in this manuscript describe two opposite extremes of a continuum, namely cytokine-dependence and cytokine-independence. However, more complex dynamics may emerge in more complex scenarios, e.g., in scenarios where leukemic cells are not fully independent of cytokines but they need less stimulation than their benign counterparts or leukemic cells are less sensitive to cytokine stimulation than healthy cells. These scenarios can be included in future versions of the presented modeling framework.

Our mathematical modeling approach allows to assign patients to different risk groups based on time evolution of their individual blast counts. This strategy is complementary to the clinically used risk stratifications which rely mostly on genetic hits. The model-based patient data analysis described in the present work provides insights into the dependence of leukemic cells on hematopoietic cytokines. The models suggest that the response of leukemic cells to cytokines has impact on disease dynamics. In the future this approach could be used to identify patients with adverse and beneficial response to exogenous cytokines and to improve risk stratification approaches.

## Acknowledgements

This work was funded by research funding from the German Research Foundation DFG (Collaborative Research Center SFB 873, Maintenance and Differentiation of Stem Cells in Development and Disease)

## Author contributions

TS, AMC, ADH designed the research, ADH contributed patient data, TS performed modeling and simulations, TS, AMC, ADH interpreted the results, TS and AMC wrote the manuscript.

